# Diaphorin induces extensive transcriptional remodeling of translation, transport, metabolism, and stress responses in *Escherichia coli*

**DOI:** 10.64898/2026.10.20.760555

**Authors:** Yuka Yasuda, Rena Takasu, Yuu Hirose, Atsushi Nakabachi

## Abstract

Diaphorin is a pederin-family polyketide produced by the obligate bacterial symbiont “*Candidatus* Profftella armatura” (Betaproteobacteria) of the Asian citrus psyllid *Diaphorina citri*, in which it accumulates at millimolar concentrations. Diaphorin suppresses the growth of many bacterial species but promotes the growth and metabolic activity of *Escherichia coli*, highlighting the importance of elucidating the mechanisms underlying these contrasting effects. Although cell-free studies show that diaphorin directly affects bacterial translation, how *E. coli* transcription changes during growth promotion remains unclear. Here, we analyzed the transcriptome of *E. coli* after 7 h of exposure to 5 mM diaphorin, when its growth-promoting effect is most pronounced. Diaphorin altered the expression of 1,278 of 4,269 analyzed genes. Functional enrichment analyses revealed strong negative enrichment of translation- and ribosome-related functions, pathway-specific changes in metabolism, and enrichment of transport-, stress-response, and quorum-sensing-related functions. Ribosomal protein genes were broadly downregulated, whereas coordinated upregulation was observed in *mdtEF*, *prpBCDE*, the Gad system, and *luxS*/*lsr* genes. These findings indicate functionally structured transcriptional remodeling during diaphorin-enhanced growth. Together with previous cell-free evidence, the pronounced translation-related changes observed here highlight bacterial translation as a major process associated with the response of *E. coli* to diaphorin. One possible interpretation of the apparently contrasting observations is that an earlier stimulation of bacterial translation is followed by compensatory downregulation of translation-related genes as growth proceeds. More broadly, these transcriptional changes provide insight into diaphorin-enhanced growth in *E. coli* and offer a basis for understanding the strikingly different effects of this symbiont-derived polyketide among bacterial species.

## Introduction

Microorganisms such as bacteria and fungi produce a wide array of secondary metabolites that mediate interactions with other organisms in their environment (Crits-Christoph *et al*., 2018; Lewis, 2020) and can also play important roles in symbiotic relationships with animal hosts (Hemmerling and Piel, 2022; Kaltenpoth *et al*., 2025).

The Asian citrus psyllid, *Diaphorina citri* Kuwayama (Hemiptera: Psylloidea), is the primary vector of “*Candidatus* Liberibacter” spp. (Alphaproteobacteria), the causative agents of citrus huanglongbing (HLB), a highly destructive and currently incurable citrus disease (Hu *et al*., 2021; Killiny, 2022). Consequently, effective control of *D. citri* is a critical priority in citrus health management. Within its body cavity, *D. citri* possesses a bacteriome, a specialized organ that houses two obligate intracellular symbionts, “*Candidatus* Carsonella ruddii” (Gammaproteobacteria) and “*Candidatus* Profftella armatura” (Betaproteobacteria) (Nakabachi *et al*., 2010; Sloan *et al*., 2014; Dan *et al*., 2017; Nakabachi and Suzaki, 2024; Song *et al*., 2025). *Carsonella* functions as a primary nutritional symbiont, supplying essential amino acids lacking in the phloem sap consumed by *D. citri* (Nakabachi *et al*., 2006; Nakabachi, Ueoka, *et al*., 2013; Nakabachi and Moran, 2022; Dittmer *et al*., 2023; Yasuda *et al*., 2024; Corretto *et al*., 2025). In contrast, *Profftella* appears to be a multifunctional symbiont that likely contributes to both host nutrition and chemical defense, an unusual feature among obligate insect symbionts (Nakabachi, Ueoka, *et al*., 2013; Nakabachi, Piel, *et al*., 2020). Its highly reduced genome (∼460 kb) allocates a substantial fraction (∼15%) to genes involved in the biosynthesis of diaphorin, a polyketide belonging to the pederin family (Nakabachi, Ueoka, *et al*., 2013). The conservation of *Profftella* and its biosynthetic genes across multiple *Diaphorina* spp. indicates an important physiological role for this compound (Nakabachi, Malenovský, *et al*., 2020; Nakabachi, Piel, *et al*., 2020).

Diaphorin accumulates at relatively high levels (approximately 2–20 mM) within *D. citri* (Nakabachi and Fujikami, 2019) and has been shown to exert inhibitory effects on diverse eukaryotic organisms (Nakabachi, Ueoka, *et al*., 2013; Nakabachi and Okamura, 2019; Yamada *et al*., 2019). In prokaryotes, diaphorin displays a distinctive dual activity, suppressing the growth of many bacterial species, including *Bacillus subtilis* (Bacilli), while enhancing the growth and metabolic activity of only a limited number of bacterial species, including *Escherichia coli* (Gammaproteobacteria) (Tanabe *et al*., 2022; Izu *et al*., 2025). These findings suggest that diaphorin functions as a chemical mediator within the *D. citri* holobiont and may influence the composition of its microbiota, including *Liberibacter* spp. (Nakabachi, Nikoh, *et al*., 2013; Hu *et al*., 2021; Killiny, 2022) and entomopathogenic bacteria used for biological control, thereby potentially contributing to HLB management strategies (Izu *et al*., 2025). Moreover, its ability to enhance bacterial growth and metabolic activity in *E. coli*, a key workhorse for industrial biotechnology, highlights its potential for improving the production efficiency of valuable compounds such as recombinant proteins and industrial enzymes (Peralta-Yahya *et al*., 2012; Rosano and Ceccarelli, 2014). However, diaphorin exhibits opposite effects on different bacterial lineages, highlighting the need to clarify its mechanism of action for potential biotechnological applications. Although *in vitro* analyses have strongly suggested that the ribosome is one of the molecular targets of diaphorin (Takasu *et al*., 2023, 2024), how *E. coli* transcription changes during diaphorin-enhanced growth remains poorly understood. To characterize this response, we performed RNA-seq analysis of *E. coli* exposed to diaphorin under conditions in which its growth-promoting effect is pronounced.

## Materials and Methods

### Preparation of diaphorin

Diaphorin was extracted and purified as previously described (Tanabe *et al*., 2022). Briefly, adult *D. citri* specimens were homogenized in methanol, and the resulting crude extract was concentrated under reduced pressure. The residue was purified by reversed-phase HPLC using a Shimadzu LC-10 system (Kyoto, Japan) equipped with an Inertsil ODS-3 C18 preparative column (GL Science, Tokyo, Japan). The diaphorin-containing fractions were collected, pooled, and dried. Purified diaphorin was re-dissolved in methanol, filtered through a 0.2 µm Minisart syringe filter (Sartorius, Göttingen, Germany), and quantified by HPLC using an Inertsil ODS-3 analytical column (GL Science). The diaphorin stock solution was stored at −30 °C until use.

### Diaphorin treatment of *E. coli*

*Escherichia coli* strain JM109 transformed with pGEM-T Easy (Promega, Madison, WI, USA) were cultured in LB medium (1% Bacto tryptone, 0.5% Bacto yeast extract, and 1% NaCl; pH 7.0) supplemented with 100 μg/mL ampicillin at 37°C for 14 h with shaking at 130 rpm for pre-cultivation. Overnight culture was inoculated into fresh medium containing either 5 mM diaphorin or no diaphorin. The concentration of diaphorin (5 mM) was selected because it falls within the physiological range in *D. citri* (Nakabachi and Fujikami, 2019) and has previously been shown to enhance the growth and metabolic activity of *E. coli* cultures (Tanabe *et al*., 2022). For each treatment, three independent cultures (160 μL each) were established in separate tubes and incubated at 37°C for 7 h. An incubation period of 7 h was selected because our previous study showed that the growth-promoting effect of 5 mM diaphorin was most pronounced at this time point, with increased cell density and larger cell size relative to untreated controls, while both control and diaphorin-treated cultures were in the logarithmic growth phase (Tanabe *et al*., 2022).

### RNA-seq library preparation and sequencing

Three independent biological replicate cultures were prepared for each treatment condition (control and 5 mM diaphorin). Following 7 h of incubation, total RNA was extracted separately from each culture using the RNeasy Protect Bacteria Mini Kit (QIAGEN, Hilden, Germany), according to the manufacturer’s instructions. Briefly, bacterial cultures were stabilized by the addition of RNAprotect Bacteria Reagent, followed by enzymatic lysis with lysozyme and Proteinase K. RNA was then purified using RNeasy spin columns, with on-column DNase digestion using the RNase-Free DNase Set (QIAGEN). RNA concentration and purity were assessed using a NanoDrop spectrophotometer (Thermo Fisher Scientific, Waltham, MA, USA) and a Qubit fluorometer (Thermo Fisher Scientific). RNA integrity was evaluated by agarose gel electrophoresis. Strand-specific RNA-seq libraries were prepared using the NEBNext Ultra II Directional RNA Library Prep Kit for Illumina in combination with the NEBNext rRNA Depletion Kit (Bacteria) (New England Biolabs, Ipswich, MA, USA) according to the manufacturer’s instructions. Libraries were sequenced on an Illumina HiSeq X platform to generate 150-bp paired-end reads.

### RNA-seq data processing and differential expression analysis

Raw paired-end reads were quality-filtered and adapter-trimmed using fastp v1.0.1 (Chen, 2025) with default parameters. Transcript abundance was quantified using Salmon v0.14.1 (Patro *et al*., 2017) against the RefSeq coding sequences (CDSs) of *E. coli* K-12 MG1655 (assembly GCF_000005845.2). Quasi-mapping was performed with automatic library type detection and the --validateMappings option enabled. Gene-level counts were imported into R v4.5.1 (R Core Team, 2025) using tximport v1.36.1 (Soneson *et al*., 2016), which imports Salmon transcript quantification results into DESeq2-compatible gene-level count matrices. Sample quality was assessed by examining expression distributions, the number of detected genes, Simple Error Ratio Estimate (SERE) (Schulze *et al*., 2012), and principal component analysis (PCA). Pairwise SERE values calculated from count data were visualized as a heatmap to assess similarity among biological replicates. PCA was performed using expression values generated by variance-stabilizing transformation (VST) with the vst function implemented in DESeq2 using the default setting (blind = TRUE). The transformed expression matrix was subjected to PCA using the prcomp function in R with centering enabled and without variance scaling. No samples were excluded from subsequent analyses. The heatmap of selected diaphorin-responsive genes was generated from VST-transformed expression values. Differential expression analysis was conducted using DESeq2 v1.48.2 (Love *et al*., 2014). Statistical significance was evaluated using the Wald test with parametric dispersion estimation. *P*-values were adjusted for multiple testing using the Benjamini–Hochberg method, and genes with an adjusted *p*-value (padj) < 0.05 were considered differentially expressed. Log_2_ fold changes were shrunk using the apeglm method for effect-size estimation and visualization, whereas Wald statistics, *p*-values, and adjusted *p*-values were obtained from the original DESeq2 model without shrinkage.

### Functional enrichment analysis

To identify biological functions associated with diaphorin-responsive genes, Gene Ontology (GO) and Kyoto Encyclopedia of Genes and Genomes (KEGG) pathway enrichment analyses were performed using goseq v1.60.0 (Young *et al*., 2010). Gene length bias inherent to RNA-seq data was corrected using the probability weighting function implemented in goseq. Functional annotations were obtained from the org.EcK12.eg.db v3.21.0 and GO.db v3.21.0 packages. KEGG pathway information for *E. coli* K-12 MG1655 (eco) was retrieved from the KEGG API using KEGGREST v1.48.1 in R. Genes detected in the RNA-seq dataset and successfully annotated with gene length information were used as the background gene set for enrichment analyses. Only GO terms and KEGG pathways containing at least two DEGs were included in the enrichment analysis. *P*-values were adjusted for multiple testing using the Benjamini–Hochberg method, and GO terms and KEGG pathways with padj < 0.05 were considered significantly enriched.

### Gene set enrichment analysis

Gene Set Enrichment Analysis (GSEA) was performed using fgsea v1.34.2 (Korotkevich *et al*., 2021), a fast preranked implementation of GSEA. All genes were ranked according to the Wald statistic from the DESeq2 differential expression analysis, with positive and negative statistics corresponding to higher and lower expression, respectively, in diaphorin-treated cells relative to controls. GO and KEGG gene sets containing between 5 and 500 genes were tested using permutation-based enrichment analysis. *P*-values were adjusted for multiple testing using the Benjamini–Hochberg method, and gene sets with padj < 0.05 were considered significantly enriched.

### Sequence data accession numbers

The raw RNA-seq data generated in this study have been deposited in the DDBJ Sequence Read Archive (DRA) under BioProject accession number PRJDB42495, with BioSample accession numbers SAMD01928879–SAMD01928884 and Run accession numbers DRR1063411–DRR1063416.

## Results

### Diaphorin-treated *E. coli* cells exhibit a distinct global transcriptional profile

Following quality filtering, 2,192,339–4,593,260 high-quality read pairs were obtained per sample, with mapping rates ranging from 66.1% to 71.8% (mean, 69.3%) (**Table S1**). Raw read counts, transcripts per million (TPM) values, and DESeq2-normalized counts for individual genes are provided in **Tables S2–S4**, respectively. Expression distributions and the numbers of detected genes were generally comparable among samples, and SERE analysis supported the overall reproducibility of the biological replicates (**Fig. S1**).

Principal component analysis (PCA) based on variance-stabilized expression values clearly separated the control and diaphorin-treated samples along PC1, which accounted for 79.5% of the total variance (**Fig. 1A**). PC2 accounted for an additional 7.9% of the variance and primarily reflected variation among the control replicates. Despite this within-group variation, all diaphorin-treated samples clustered separately from the controls, indicating that diaphorin treatment was associated with a pronounced global shift in the *E. coli* transcriptome.

**Figure 1.**
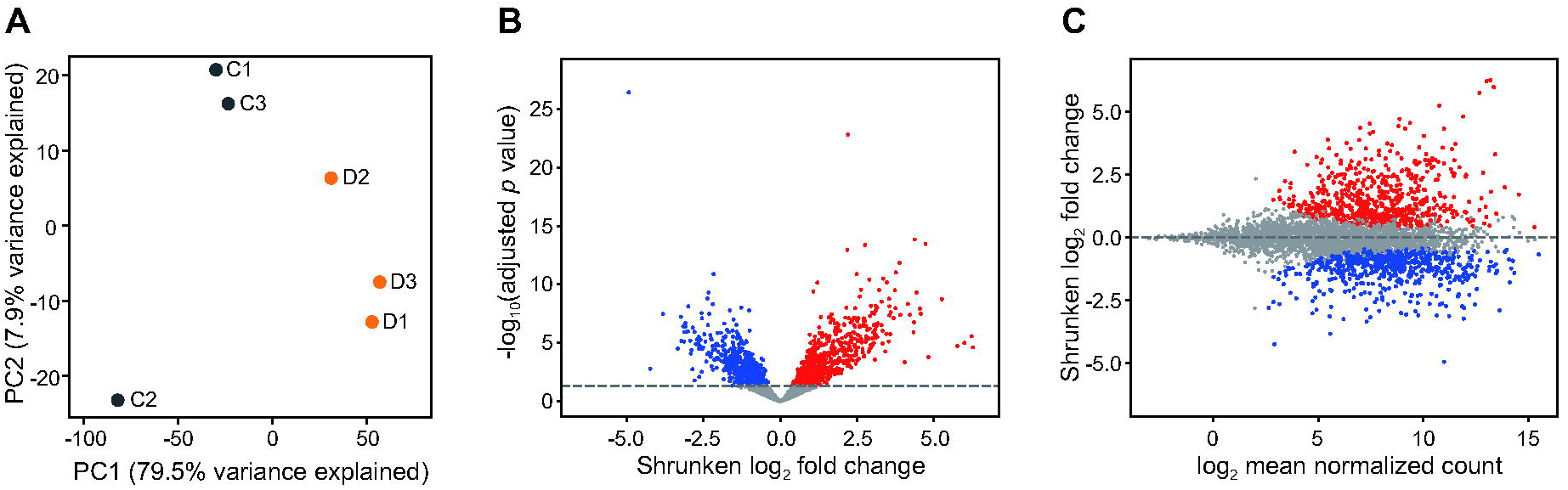
Global transcriptomic response of *E. coli* to diaphorin treatment. (A) Principal component analysis of RNA-seq samples. Principal component analysis (PCA) was performed using variance-stabilized expression values generated by DESeq2. Each point represents one biological replicate (Control, *n* = 3; Diaphorin-treated, *n* = 3). Percentages on the axes indicate the proportion of total variance explained by each principal component. (B) Volcano plot of differential gene expression. The x-axis represents apeglm-shrunken log_2_ fold changes (diaphorin-treated vs. control), and the y-axis represents −log_10_(padj). Red and blue points indicate significantly upregulated and downregulated genes, respectively (padj < 0.05), whereas grey points indicate non-significant genes. The dashed horizontal line indicates padj = 0.05. (C) MA plot of differential gene expression. The x-axis represents the log_2_-transformed mean DESeq2-normalized count across all samples, and the y-axis represents the apeglm-shrunken log_2_ fold change. Red and blue points indicate significantly upregulated and downregulated genes, respectively (padj < 0.05), whereas grey points indicate non-significant genes. The dashed horizontal line indicates a log_2_ fold change of zero.

### Diaphorin treatment affects the expression of more than 1,200 *E. coli* genes

Differential expression analysis identified 1,278 of the 4,269 analyzed genes (29.9%) as significantly differentially expressed following diaphorin treatment (padj < 0.05), comprising 635 upregulated and 643 downregulated genes. Thus, the transcriptional response involved nearly one-third of the analyzed genes, with nearly equal numbers of genes showing increased and decreased expression.

The volcano plot showed significant expression changes in both directions over a wide range of fold changes (**Fig. 1B**), whereas the MA plot showed that differentially expressed genes were distributed across a broad range of mean expression levels (**Fig. 1C**). Complete expression and differential-expression statistics for all 4,269 genes are provided in **Table S5**.

### Functional enrichment reveals extensive transcriptional changes across multiple cellular functions

GO enrichment analysis of the DEGs revealed prominent enrichment of functions associated with protein synthesis (**Fig. 2**; **Table S7**). Among Biological Process terms, translation (GO:0006412) and cytoplasmic translation (GO:0002181) were among the 20 terms with the smallest adjusted *p*-values, with translation containing the largest number of DEGs in this set (**Fig. 2**). Enrichment of translation-related functions was further reflected in terms associated with ribosomal small- and large-subunit assembly, tRNA processing, and tRNA and rRNA binding.

**Figure 2.**
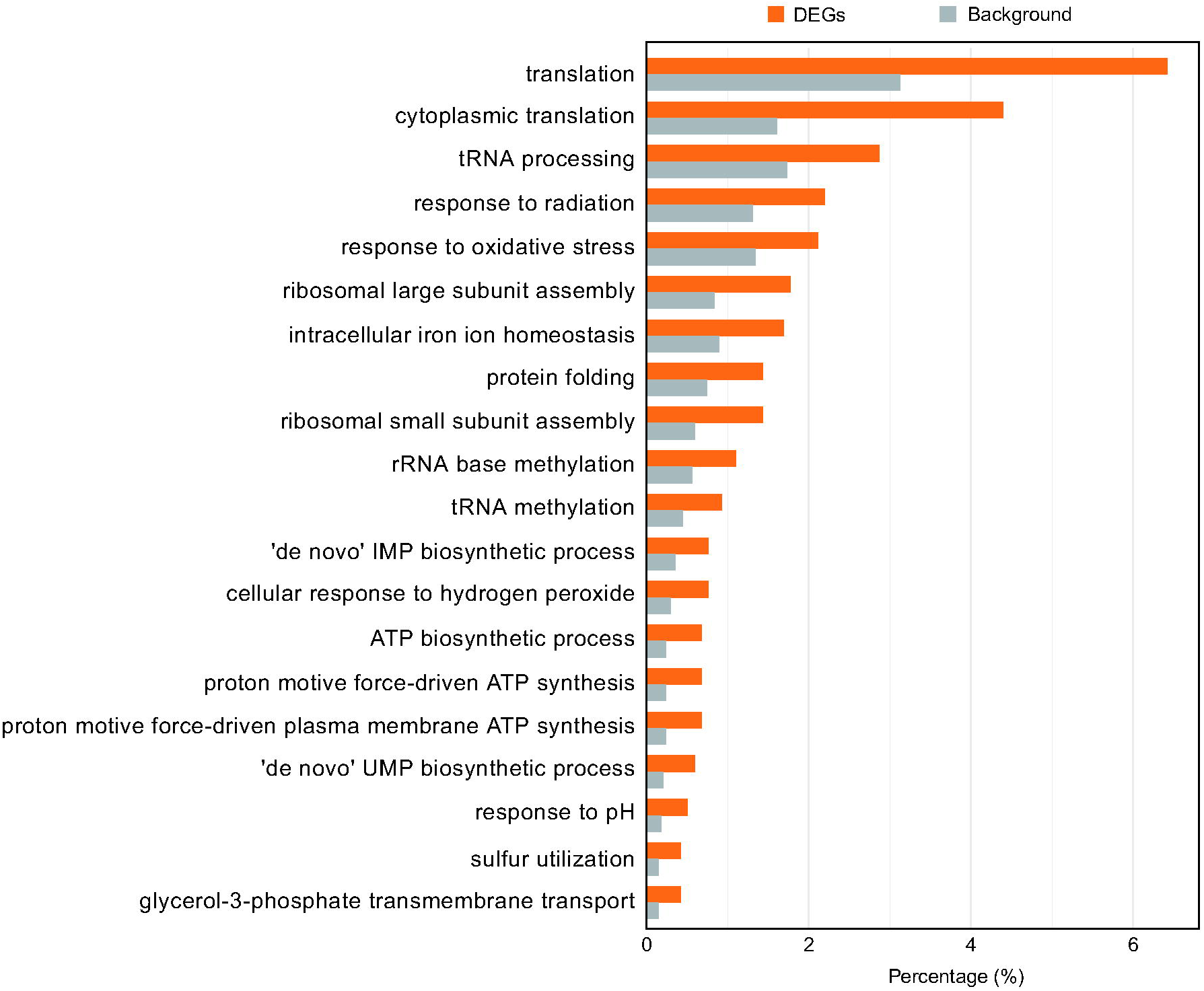
GO enrichment analysis of differentially expressed genes (DEGs). Double bar plot showing the top 20 significantly enriched Gene Ontology (GO) terms in the Biological Process (BP) category. The x-axis represents the percentage of genes assigned to each GO term. Orange bars (DEGs) indicate the percentage of genes in each term relative to the total number of GO-annotated DEGs (n = 1,183), whereas gray bars (Background) indicate the percentage relative to the total number of GO-annotated genes included in the enrichment analysis (n = 3,354). The 20 terms with the smallest adjusted *p*-values (padj) are shown.

Beyond protein synthesis, enriched GO terms included ATP biosynthetic process and proton motive force-driven ATP synthesis, as well as functions related to cellular stress and homeostasis, including intracellular iron ion homeostasis, cellular response to hydrogen peroxide, and protein folding. The de novo UMP biosynthetic process was also enriched. In the KEGG over-representation analysis, Ribosome (eco03010) was the only significantly enriched pathway (**Table S8**).

GSEA further revealed the direction of coordinated transcriptional changes across gene sets. A total of 115 GO terms and 19 KEGG pathways were significantly enriched in response to diaphorin treatment (padj < 0.05; **Tables S9** and **S10**). GO and KEGG GSEA showed concordant enrichment patterns across several major cellular functions. Translation- and ribosome-related gene sets showed particularly strong negative enrichment, including the GO term cytoplasmic translation (normalized enrichment score [NES] = −3.29, padj = 3.67 × 10 ²) and the KEGG Ribosome pathway (NES = −3.20, padj = 9.29 × 10 ²; **Fig. 3A**). Negative enrichment extended to aminoacyl-tRNA-related functions in both analyses.

**Figure 3.**
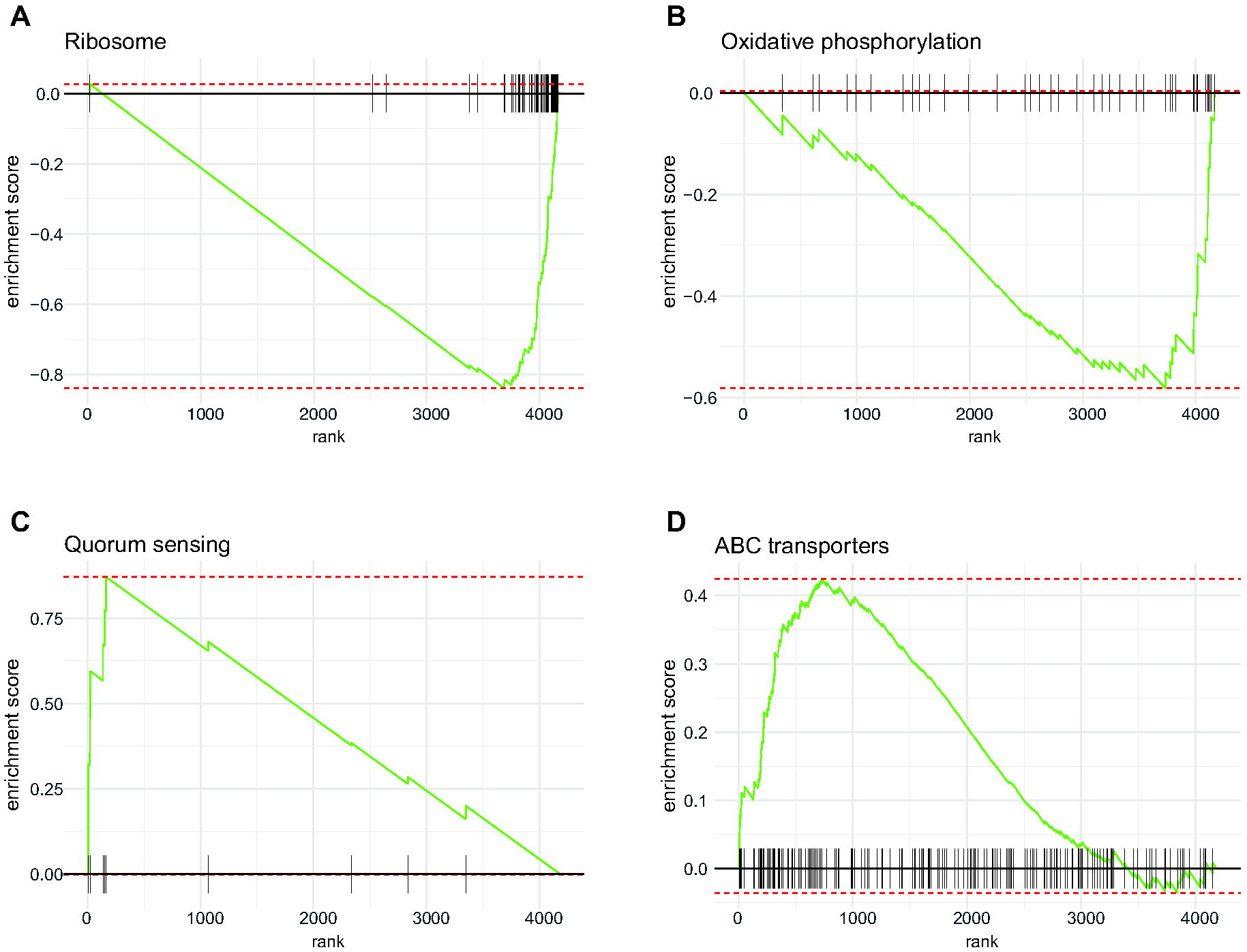
Gene set enrichment analysis (GSEA) of representative gene sets affected by diaphorin treatment. GSEA plots for (A) the KEGG Ribosome pathway, (B) the KEGG Oxidative phosphorylation pathway, (C) the GO term quorum sensing, and (D) the KEGG ABC transporters pathway. Genes were ranked according to the Wald statistic from DESeq2 differential-expression analysis, with positive values corresponding to higher expression in diaphorin-treated cells relative to controls. The green line indicates the running enrichment score, and black vertical lines indicate the positions of genes belonging to each pathway in the ranked gene list.

Concordant negative enrichment was also observed for nucleotide biosynthesis and respiratory energy metabolism. GO terms related to purine and pyrimidine nucleotide biosynthesis were negatively enriched, consistent with negative enrichment of the KEGG Purine metabolism (NES = −1.82, padj = 9.84 × 10) and Pyrimidine metabolism (NES = −2.37, padj = 7.42 × 10) pathways. Similarly, negative enrichment of GO terms related to ATP biosynthesis and electron transport was consistent with negative enrichment of KEGG Oxidative phosphorylation (NES = −2.04, padj = 2.65 × 10; **Fig. 3B**) and the TCA cycle (NES = −1.76, padj = 1.67 × 10 ²).

In contrast, carbohydrate utilization showed concordant positive enrichment in both analyses. The GO terms glycolytic process and phosphoenolpyruvate-dependent sugar phosphotransferase system were positively enriched, consistent with positive enrichment of the KEGG Glycolysis / Gluconeogenesis (NES = 1.65, padj = 2.83 × 10 ²) and Phosphotransferase system (PTS) (NES = 1.87, padj = 4.61 × 10 ³) pathways. Other carbohydrate-related pathways, including Starch and sucrose metabolism (NES = 2.14, padj = 2.65 × 10) and Fructose and mannose metabolism (NES = 2.01, padj = 5.46 × 10), were also positively enriched. Fatty acid metabolism showed a similarly contrasting pattern, with negative enrichment of Fatty acid biosynthesis (NES = −1.97, padj = 1.37 × 10 ³) and positive enrichment of Fatty acid degradation (NES = 1.89, padj = 1.67 × 10 ²), the latter being consistent with positive enrichment of the GO term fatty acid beta-oxidation.

Positive enrichment was also observed for the GO term quorum sensing (NES = 2.13, padj = 1.32 × 10; **Fig. 3C**) and the KEGG ABC transporters pathway (NES = 1.80, padj = 1.05 × 10; Fig. 3d). ABC transporter-related gene sets were likewise positively enriched in the GO analysis (NES = 2.22, padj = 3.71 × 10). The GO term autoinducer AI-2 transmembrane transport was also positively enriched.

### Gene-level analysis identifies coordinated responses in acid resistance, efflux, quorum sensing, propionate catabolism, and ribosomal proteins

Examination of individual differentially expressed genes further identified prominent coordinated responses to diaphorin in acid resistance, multidrug efflux, quorum sensing, propionate catabolism, and ribosomal proteins (**Fig. 4**; **Table S6**). Among the acid-resistance genes, the glutamate-dependent acid-resistance (Gad) system showed coordinated upregulation: the core components *gadA*, *gadB*, and *gadC* and the regulatory genes *gadE*, *gadX*, and *gadW* were significantly upregulated (padj < 0.05). Particularly strong induction was observed for *gadA*, *gadB*, *gadC*, and *gadE*, with shrunken log2 fold changes of 4.54–5.26. The genes encoding the acid-stress-responsive periplasmic chaperones HdeA and HdeB were also significantly upregulated, with shrunken log_2_ fold changes of 4.34 and 4.05, respectively.

**Figure 4.**
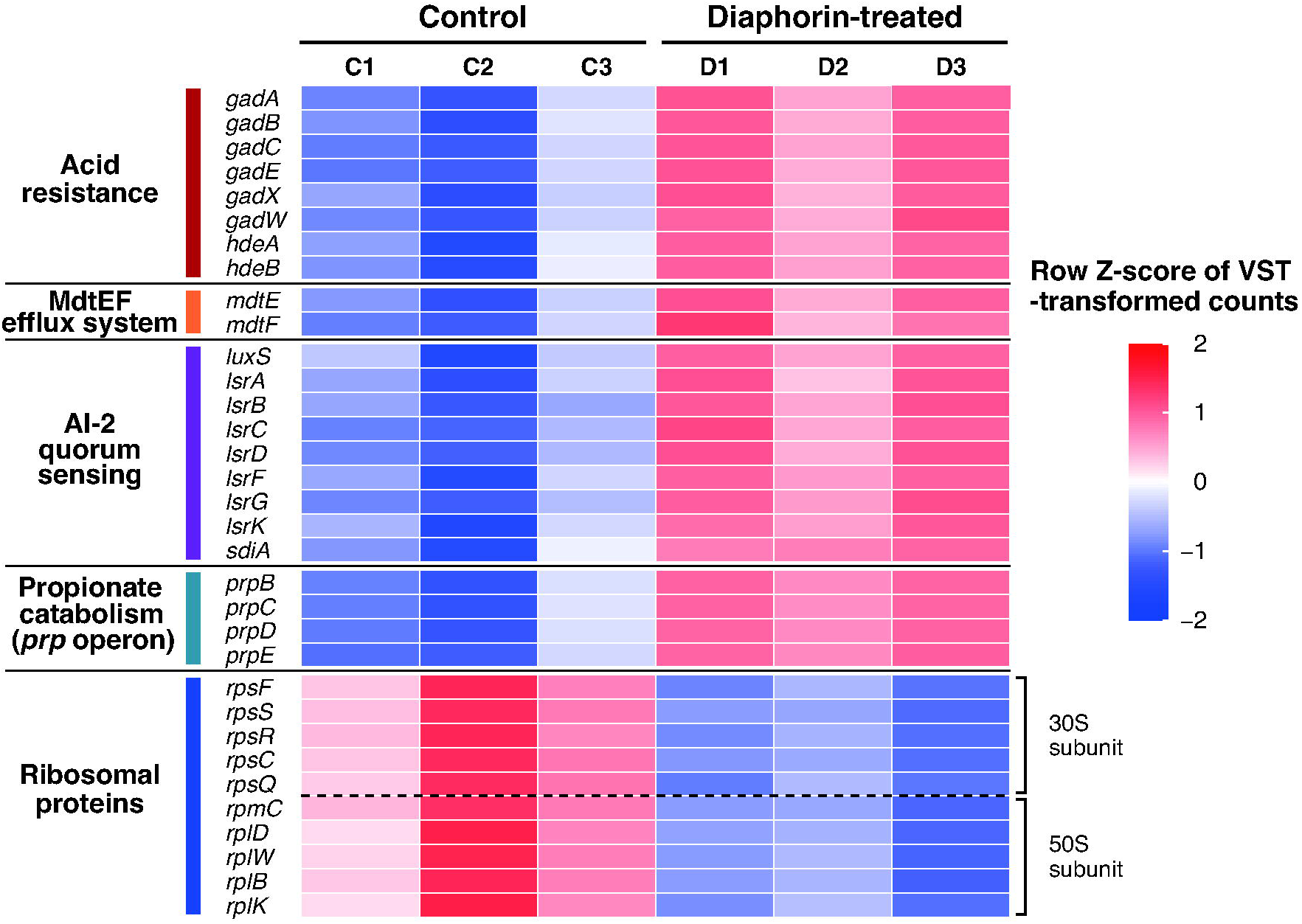
Representative transcriptional modules responsive to diaphorin. Heatmap of selected genes representing major biological processes affected by diaphorin treatment. Genes were selected to represent major biological processes highlighted by differential-expression and functional enrichment analyses, including acid resistance (the *gad* regulon and acid-stress genes), MdtEF efflux system, AI-2 quorum sensing (luxS/lsr operon), propionate catabolism (*prp* operon), and ribosomal proteins. Expression values were variance-stabilized using the DESeq2 variance stabilizing transformation (VST) and row-scaled (Z-score). Red indicates higher expression relative to the row mean, and blue indicates lower expression. Samples: C1–C3, control; D1–D3, diaphorin-treated.

Both components of the MdtEF multidrug efflux system were significantly upregulated, with shrunken log2 fold changes of 2.80 for *mdtE* and 1.98 for *mdtF*. A coordinated response was also evident in the autoinducer-2 (AI-2) quorum-sensing system.

The AI-2 production gene *luxS* and the *lsr* genes involved in AI-2 uptake, phosphorylation, and processing (*lsrA*–*D*, *lsrF*, *lsrG*, and *lsrK*) were significantly upregulated. The *lsr* genes generally showed larger expression changes than *luxS*, with several exhibiting shrunken log2 fold changes greater than 3.0. The quorum-sensing-related regulatory gene *sdiA* was also significantly upregulated. This coordinated gene-level response was consistent with the positive enrichment of the GO term quorum sensing and autoinducer AI-2 transmembrane transport observed by GSEA (**Fig. 3C**).

A particularly strong coordinated increase was observed in propionate catabolism. All four genes of the *prpBCDE* operon were strongly upregulated, with shrunken log2 fold changes ranging from 5.76 to 6.27 (**Fig. 4**; **Table S6**). This represented one of the most pronounced coordinated gene-level responses identified.

In contrast to the predominantly upregulated modules described above, ribosomal protein genes showed a broad and highly consistent decrease in expression, encompassing genes encoding proteins of both the 30S and 50S ribosomal subunits. All 54 ribosomal protein genes showed negative shrunken log2 fold changes (range, −3.12 to −0.55; median, −1.64), with 52 reaching statistical significance (padj < 0.05). Representative genes illustrating this coordinated decrease are shown in **Fig. 4**. This broad decrease was consistent with the strong negative enrichment of translation- and ribosome-related gene sets observed by GSEA (**Fig. 3A**).

## Discussion

In this study, we characterized the transcriptomic state of *E. coli* after continuous exposure to diaphorin for 7 h. This time point corresponds to the logarithmic growth phase and, importantly, to the point at which the growth-promoting effect of 5 mM diaphorin was most pronounced in our previous study, with the ratio of ΔOD600 between diaphorin-treated and control cultures reaching a maximum of 1.40. At this time point, diaphorin-treated cultures also contained 12.3% more cells, and the cells had a 29.4% larger mean volume than control cells. In addition, diaphorin increased

β-galactosidase activity, indicating enhanced metabolic activity (Tanabe *et al*., 2022). Thus, the present RNA-seq analysis was designed to characterize the transcriptional state associated with the established growth-promoting phenotype rather than the immediate transcriptional response to diaphorin exposure. Diaphorin treatment was associated with extensive transcriptomic changes, with approximately 30% of the analyzed genes being differentially expressed. Functional analyses revealed broad changes in translation, transport, stress responses, quorum sensing, and multiple metabolic processes, with contrasting transcriptional responses among pathways of carbon, energy, nucleotide, and fatty acid metabolism. At the gene level, ribosomal protein genes were broadly downregulated, whereas *mdtE* and *mdtF*, the *prpBCDE* operon, genes of the glutamate-dependent acid resistance system, and genes associated with AI-2 signaling were upregulated. Together, these findings characterize a distinct transcriptional state associated with diaphorin-enhanced growth, likely reflecting both responses to diaphorin itself and secondary adjustments accompanying the altered physiological state.

The pronounced alteration of translation- and ribosome-related functions was particularly notable. Despite the transcriptome being sampled at the time point at which the diaphorin-associated growth advantage was maximal, all 54 ribosomal protein genes showed negative shrunken log2 fold changes, and translation- and ribosome-related gene sets showed some of the strongest negative enrichment. Negative enrichment was also evident in aminoacyl-tRNA-related functions in both GO and KEGG GSEA, indicating that the response extended beyond ribosomal components to another major component of the translation machinery. This broad translation-related response is especially intriguing in light of previous evidence that bacterial translation is affected by diaphorin.

Pederin-family compounds are known to inhibit eukaryotic protein synthesis through interaction with the ribosome (Tiboni *et al*., 1968; Dmitriev *et al*., 2020), whereas much less was initially known about their effects on bacterial ribosomes. Subsequent cell-free studies demonstrated concentration-dependent effects of diaphorin on bacterial protein synthesis, with inhibition at relatively high concentrations but enhancement within a limited lower concentration range (Takasu *et al*., 2023, 2024). These direct effects raise the possibility that the bacterial translational machinery is among the major cellular targets of diaphorin. From this perspective, the strong downregulation of translation-related functions observed here need not reflect a direct inhibitory effect of diaphorin on the translational machinery. One possibility is that enhanced translational activity earlier during diaphorin exposure may have triggered compensatory regulation that reduced translation-related gene expression by the 7-h mid-log phase examined here. Such a scenario could provide a plausible explanation for the otherwise paradoxical combination of enhanced growth-related phenotypes and widespread downregulation of translation-related genes.

Transport-related transcriptional changes were also prominent. The multidrug efflux genes *mdtE* and *mdtF* were upregulated in diaphorin-treated cells, and ABC transporter-related functions were positively enriched in both KEGG and GO GSEA (**Fig. 3D**), indicating broader transcriptional changes in transport functions beyond the MdtEF system. MdtEF is a resistance-nodulation-cell division (RND)-type multidrug efflux system that functions with the outer-membrane channel TolC and confers resistance to a range of structurally diverse compounds in *E. coli* (Nishino and Yamaguchi, 2002). Although the substrates of MdtEF characterized to date do not include diaphorin, its broad substrate range raises the intriguing possibility that increased efflux capacity could reduce intracellular accumulation of diaphorin and thereby contribute to the ability of *E. coli* to tolerate or benefit from exposure to the compound. This possibility is particularly interesting in light of the strikingly different effects of diaphorin among bacterial species. Our previous study showed that 5 mM diaphorin promoted growth and metabolic activity of *E. coli* but strongly inhibited growth and cell division of *B. subtilis* (Tanabe *et al*., 2022). Subsequent studies further revealed that the response cannot be explained simply by differences in cell-envelope architecture between

Gram-positive and Gram-negative bacteria, because growth promotion was observed in some Gram-positive bacteria, whereas growth inhibition occurred in many Gram-negative species (Izu *et al*., 2025). Thus, susceptibility to diaphorin appears to depend on species-specific cellular properties. Differences in the repertoire, regulation, and activity of transport systems could be among the factors determining intracellular exposure to diaphorin or associated metabolites and thereby contribute to these species-dependent responses.

Metabolic gene expression also showed extensive pathway-specific changes. GO and KEGG GSEA showed concordant enrichment patterns for several major aspects of metabolism. Rather than showing a uniform directional response, central carbon and energy metabolism exhibited contrasting changes: the TCA cycle and oxidative phosphorylation were negatively enriched, whereas glycolysis/gluconeogenesis and the PTS were positively enriched. Other carbohydrate-related pathways, including fructose/mannose and starch/sucrose metabolism, were likewise positively enriched. Lipid metabolism showed a similarly contrasting pattern, with fatty acid biosynthesis negatively enriched and fatty acid degradation positively enriched. Together, these patterns indicate extensive but selective reorganization of metabolic gene expression at the 7-h time point rather than generalized metabolic activation or suppression.

Within this broader metabolic response, particularly strong induction was observed for the *prpBCDE* operon. The *prp* genes encode the 2-methylcitrate pathway for propionate catabolism, in which propionyl-CoA is converted through 2-methylcitrate intermediates to pyruvate and succinate (Brock *et al*., 2002). Because diaphorin-treated and control cells were grown in the same LB medium, the marked differential induction of this pathway points to a diaphorin-associated change in the production, availability, or utilization of metabolites feeding into propionate/propionyl-CoA metabolism rather than simply reflecting differences in medium composition.

The striking induction of the *prp* pathway may also be relevant to another prominent transcriptional response observed here: activation of the glutamate-dependent acid resistance system. Recent work has shown that short-chain fatty acids, including propionate, strongly induce the Gad system in *E. coli*, with effects that cannot be explained solely by the accompanying decrease in extracellular pH (Guo *et al*., 2026). In the present study, *gadA*, *gadB*, *gadC*, and *gadE* were among the most strongly induced genes, accompanied by increased expression of *gadX*, *gadW*, *hdeA*, and *hdeB*. The simultaneous strong induction of the *prp* and Gad systems therefore raises the possibility that altered propionate metabolism contributes to activation of the acid-resistance network during growth in the presence of diaphorin. This potential metabolic link suggests that the pronounced *prp* and Gad responses may represent interconnected components of the diaphorin-associated physiological state. The Gad response may also be linked to the induction of *mdtEF*: in *E. coli*, *mdtEF* is co-transcribed with *gadE* as part of the *gadE-mdtEF* operon (Hirakawa *et al*., 2006), and GadX has been shown to activate *mdtEF* expression (Nishino *et al*., 2008). Thus, the transcriptional induction of *mdtEF* may also arise as a component of the broader Gad-associated response.

Diaphorin treatment also affected the AI-2 quorum-sensing system. In *E. coli*, LuxS is involved in the production of autoinducer-2 (AI-2), a signaling molecule associated with population-density-dependent regulation, while the Lsr system mediates AI-2 uptake and intracellular processing (Xavier and Bassler, 2005). The coordinated upregulation of *luxS* and multiple *lsr* genes indicates transcriptional activation of the AI-2 system, consistent with the positive enrichment of the GO term quorum sensing and autoinducer AI-2 transmembrane transport in GSEA. These changes suggest enhanced engagement of the AI-2 system in the physiological state established during growth in the presence of diaphorin. This response is particularly relevant at the 7-h time point, when diaphorin-treated cultures contained more cells than controls (Tanabe *et al*., 2022). Activation of the AI-2 system may therefore reflect, or contribute to, the altered population-level state associated with diaphorin-enhanced growth and metabolism.

Taken together, these findings provide a framework for understanding the unusual growth-promoting action of diaphorin on *E. coli*. One possible interpretation is that diaphorin initially stimulates protein synthesis through its effects on the bacterial translational machinery. As growth proceeds, compensatory regulation of translation-related gene expression may become apparent, accompanied by transcriptional changes in transport systems, selective metabolic reorganization, stress-response functions, and population-density-associated signaling. The transcriptional profile observed at 7 h may therefore represent a physiological state established during diaphorin-enhanced growth, resulting from both the primary cellular effects of the compound and subsequent physiological adjustments. This model may also help explain the species-dependent effects of diaphorin. Diaphorin occurs at millimolar concentrations in its natural host, *D. citri* (Nakabachi and Fujikami, 2019), and its effects on bacterial growth vary substantially among species in a manner that does not simply follow the Gram-positive/Gram-negative distinction (Tanabe *et al*., 2022; Izu *et al*., 2025). Species-specific differences in transport capacity, cellular targets, metabolic state, and regulatory responses may collectively determine whether exposure to diaphorin results in growth promotion or inhibition.

The present study examined a single diaphorin concentration at a single time point and therefore provides a snapshot of the transcriptional state associated with diaphorin-enhanced growth. In particular, the proposed temporal relationship between earlier effects on translation and the subsequent downregulation of translation-related genes remains to be tested directly. Nevertheless, the extensive and coordinated changes identified here reveal previously unrecognized features of the transcriptional response to diaphorin and provide a basis for understanding its unusual growth-promoting effect on *E. coli*.

