## Supplementary Tables S1 and S6 for "Diaphorin induces extensive transcriptional remodeling of translation, transport, metabolism, and stress responses in *Escherichia coli*"

**Supplementary Materials**

**List of supplementary materials**

Supplementary Table S1. Summary of total read pairs post-trimming and reference mapping rates for each sample.

Supplementary Table S2. Raw read counts for each sample (provided as a separate Excel file).

Supplementary Table S3. Transcripts per million (TPM) values for each sample (provided as a separate Excel file).

Supplementary Table S4. DESeq2-normalized read counts for each sample (provided as a separate Excel file).

Supplementary Table S5. Expression data and DESeq2 differential expression results for 4,269 *Escherichia coli* genes following diaphorin exposure (provided as a separate Excel file).

Supplementary Table S6. Differential expression of representative genes shown in Fig. 4.

Supplementary Table S7. Results of Gene Ontology (GO) over-representation analysis (provided as a separate Excel file).

Supplementary Table S8. Results of KEGG pathway over-representation analysis (provided as a separate Excel file).

Supplementary Table S9. GSEA results for GO terms (provided as a separate Excel file).

Supplementary Table S10. GSEA results for KEGG pathways (provided as a separate Excel file).

Supplementary Fig. S1. Quality assessment of RNA-seq data.

**Supplementary Table S1.** Summary of total read pairs post-trimming and reference mapping rates for each sample.

| **Sample** | **Total reads** | **Mapping rate (%)** |
| --- | --- | --- |
| Ecoli-C1 | 4,368,087 | 71.8 |
| Ecoli-C2 | 4,593,260 | 66.1 |
| Ecoli-C3 | 3,895,622 | 66.3 |
| Ecoli-D1 | 2,808,674 | 70.8 |
| Ecoli-D2 | 3,170,217 | 71.5 |
| Ecoli-D3 | 2,192,339 | 69.7 |

**Supplementary Table S6.** Differential expression of representative genes shown in Fig. 4.

| **Functional module** | **Gene** | **Overall mean normalized count** | **Mean normalized count, control** | **Mean normalized count, diaphorin** | **log_2_FC** | **Shrunken**  **log_2_FC** | **lfcSE** | **Wald statistic** | ***p*-value** | **Adjusted**  ***p*-value** |
| --- | --- | --- | --- | --- | --- | --- | --- | --- | --- | --- |
| Acid resistance | *gadA* | 659.14 | 45.27 | 1273.02 | 4.817 | 4.571 | 0.794 | 6.248 | 4.15E-10 | 3.34E-08 |
| Acid resistance | *gadB* | 3821.64 | 168.97 | 7474.32 | 5.467 | 4.824 | 1.395 | 4.299 | 1.72E-05 | 1.71E-04 |
| Acid resistance | *gadC* | 1746.22 | 75.83 | 3416.61 | 5.496 | 5.257 | 0.831 | 6.779 | 1.21E-11 | 1.87E-09 |
| Acid resistance | *gadE* | 175.44 | 12.52 | 338.37 | 4.771 | 4.539 | 0.762 | 6.437 | 1.22E-10 | 1.22E-08 |
| Acid resistance | *gadX* | 165.99 | 40.11 | 291.87 | 2.864 | 2.701 | 0.511 | 5.760 | 8.43E-09 | 3.46E-07 |
| Acid resistance | *gadW* | 75.03 | 22.42 | 127.64 | 2.513 | 2.358 | 0.470 | 5.488 | 4.08E-08 | 1.25E-06 |
| Acid resistance | *hdeA* | 2029.90 | 155.51 | 3904.29 | 4.651 | 4.338 | 0.883 | 5.481 | 4.24E-08 | 1.28E-06 |
| Acid resistance | *hdeB* | 1046.08 | 78.29 | 2013.87 | 4.686 | 4.048 | 1.292 | 4.020 | 5.83E-05 | 4.48E-04 |
| MdtEF efflux system | *mdtE* | 43.99 | 9.79 | 78.19 | 3.004 | 2.799 | 0.584 | 5.314 | 1.07E-07 | 2.73E-06 |
| MdtEF efflux system | *mdtF* | 104.21 | 38.60 | 169.82 | 2.142 | 1.980 | 0.454 | 4.860 | 1.17E-06 | 1.86E-05 |
| AI-2 quorum sensing | *luxS* | 791.45 | 435.24 | 1147.66 | 1.398 | 1.314 | 0.290 | 4.875 | 1.09E-06 | 1.74E-05 |
| AI-2 quorum sensing | *lsrA* | 93.82 | 20.84 | 166.80 | 3.002 | 2.773 | 0.617 | 5.053 | 4.35E-07 | 8.29E-06 |
| AI-2 quorum sensing | *lsrB* | 123.56 | 14.39 | 232.72 | 4.011 | 3.886 | 0.515 | 7.903 | 2.72E-15 | 1.44E-12 |
| AI-2 quorum sensing | *lsrC* | 58.19 | 9.85 | 106.52 | 3.446 | 3.298 | 0.525 | 6.704 | 2.03E-11 | 3.00E-09 |
| AI-2 quorum sensing | *lsrD* | 45.43 | 6.47 | 84.40 | 3.718 | 3.548 | 0.581 | 6.535 | 6.37E-11 | 7.60E-09 |
| AI-2 quorum sensing | *lsrF* | 152.71 | 27.80 | 277.63 | 3.322 | 3.126 | 0.596 | 5.743 | 9.32E-09 | 3.70E-07 |
| AI-2 quorum sensing | *lsrG* | 81.44 | 10.33 | 152.56 | 3.895 | 3.766 | 0.518 | 7.650 | 2.00E-14 | 9.26E-12 |
| AI-2 quorum sensing | *lsrK* | 159.83 | 60.25 | 259.41 | 2.105 | 1.953 | 0.439 | 4.933 | 8.10E-07 | 1.37E-05 |
| AI-2 quorum sensing | *sdiA* | 118.43 | 51.16 | 185.70 | 1.863 | 1.720 | 0.408 | 4.687 | 2.77E-06 | 3.79E-05 |
| Propionate catabolism (*prp* operon) | *prpB* | 9410.53 | 155.51 | 18665.55 | 6.908 | 6.270 | 1.552 | 4.798 | 1.61E-06 | 2.44E-05 |
| Propionate catabolism (*prp* operon) | *prpC* | 6577.40 | 161.94 | 12992.86 | 6.327 | 5.761 | 1.393 | 4.858 | 1.18E-06 | 1.86E-05 |
| Propionate catabolism (*prp* operon) | *prpD* | 10537.58 | 224.47 | 20850.68 | 6.538 | 5.991 | 1.392 | 5.001 | 5.71E-07 | 1.03E-05 |
| Propionate catabolism (*prp* operon) | *prpE* | 8278.95 | 155.72 | 16402.18 | 6.719 | 6.227 | 1.332 | 5.318 | 1.05E-07 | 2.70E-06 |
| Ribosomal proteins (30S subunit) | *rpsF* | 2402.11 | 4373.09 | 431.12 | -3.342 | -3.118 | 0.628 | -5.496 | 3.88E-08 | 1.21E-06 |
| Ribosomal proteins (30S subunit) | *rpsS* | 1590.06 | 2886.10 | 294.02 | -3.294 | -3.092 | 0.593 | -5.716 | 1.09E-08 | 4.24E-07 |
| Ribosomal proteins (30S subunit) | *rpsR* | 581.66 | 1054.27 | 109.05 | -3.271 | -3.055 | 0.613 | -5.511 | 3.57E-08 | 1.15E-06 |
| Ribosomal proteins (30S subunit) | *rpsC* | 3956.48 | 7027.96 | 884.99 | -2.989 | -2.794 | 0.559 | -5.507 | 3.66E-08 | 1.17E-06 |
| Ribosomal proteins (30S subunit) | *rpsQ* | 853.58 | 1493.50 | 213.67 | -2.804 | -2.599 | 0.562 | -5.161 | 2.45E-07 | 5.22E-06 |
| Ribosomal proteins (50S subunit) | *rpmC* | 869.75 | 1577.30 | 162.19 | -3.280 | -3.091 | 0.573 | -5.877 | 4.17E-09 | 2.08E-07 |
| Ribosomal proteins (50S subunit) | *rplD* | 3220.73 | 5519.24 | 922.22 | -2.581 | -2.333 | 0.596 | -4.519 | 6.21E-06 | 7.27E-05 |
| Ribosomal proteins (50S subunit) | *rplW* | 1908.58 | 3352.18 | 464.97 | -2.849 | -2.609 | 0.611 | -4.852 | 1.22E-06 | 1.91E-05 |
| Ribosomal proteins (50S subunit) | *rplB* | 5631.44 | 9915.32 | 1347.55 | -2.879 | -2.655 | 0.591 | -5.052 | 4.38E-07 | 8.29E-06 |
| Ribosomal proteins (50S subunit) | *rplK* | 1578.34 | 2703.69 | 453.00 | -2.577 | -2.320 | 0.607 | -4.440 | 9.01E-06 | 9.91E-05 |

*Genes are ordered as displayed in Fig. 4. Statistical values were extracted from Table S5 (DESeq2 differential-expression analysis); shrunk_log2FC denotes the apeglm-shrunken log2 fold change.*

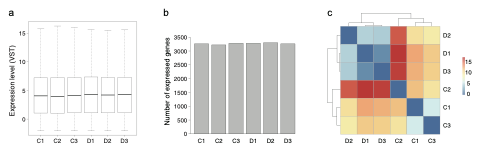

**Supplementary Fig. S1.** Quality assessment of RNA-seq data. (a) Distribution of variance-stabilizing transformed (VST) expression values across samples. (b) Number of expressed genes detected in each sample. (c) Heatmap of SERE distances among samples with hierarchical clustering. C1–C3, control samples; D1–D3, diaphorin-treated samples.
